# The effect of sword bean extract on the relationship between rheumatoid arthritis and periodontitis in mice

**DOI:** 10.1101/546788

**Authors:** Kosei Matsumoto, Yuko Nakatsuka, Kaname Shirai, Shintaro Shimizu, Shunshuke Yanase, Yoshihiro Abiko, Yasushi Furuichi

**Affiliations:** Division of Periodontology and Endodontology Department of Oral Rehabilitation School of Dentistry, Health Sciences University of Hokkaido, Tobetsu, Hokkaido, Japan; Division of Oral Medicine and Pathology, School of Dentistry, Health Sciences University of Hokkaido, Tobetsu, Hokkaido, Japan

## Abstract

**Objectives:** Several studies in humans and experimental animals have reported an interaction between rheumatoid arthritis (RA) and periodontitis (PD). We previously showed that extracts of *Canavalia gladiata* (sword bean extract, SBE) can treat PD in rats. Here, we investigated the relationship between RA and PD and the effects of SBE in an experimental mouse model.

**Methods:** Female SKG mice were assigned to eight groups (n=6/group): (1) Untreated controls, (2) RA (induced at 6 weeks of age), (3) PD (induced at 10 weeks of age), (4) RA + PD, (5) SBE (2 mg/ml in drinking water starting at 5 weeks of age), (6) PD + SBE, (7) RA + SBE, and (8) RA + PD + SBE. Mice were sacrificed at 13 weeks of age, and alveolar bone resorption, periodontal tissue inflammation, and paw joint inflammation were assessed by histology and immunohistochemistry.

**Results:** Mice in the RA + PD group exhibited significantly higher inflammation scores in the joint tissues as well as more abundant IL-17-positive cells and cathepsin K-positive osteoclasts in the radial bone compared with the RA mice. Alveolar bone resorption was also significantly more severe in the RA + PD mice than in the PD mice. SBE treatment significantly improved all bone resorption and tissue inflammation scores in mice with RA + PD.

**Conclusion:** Concomitant RA and PD exacerbates the tissue destruction symptomatic of each condition. SBE suppresses all parameters evaluated, suggesting that it is has anti-inflammatory activities in both RA and PD.

## Introduction

Periodontitis (PD) is caused by infection of periodontal tissue with oral bacterial communities and is the most common chronic inflammatory disease affecting humans [1]. The disease is characterized by destruction of connective tissue and alveolar bone, both of which are fundamental to tooth support. Recent work has suggested that PD can influence systemic diseases and conditions such as diabetes, arteriosclerosis, and rheumatoid arthritis (RA), and, in pregnant women, can result in low birth weight in the offspring [2]. RA is a systemic autoimmune disease characterized by chronic inflammation of the joints, leading to bone erosion and progressive disability. Both RA and PD are associated with inflammatory cell infiltration that mediates the tissue destruction [3, 4].

Several epidemiological studies have evaluated the association between RA and PD. Some studies showed a significantly higher incidence of tooth loss and alveolar bone loss in patients with RA [5, 6], and one study reported that PD is a risk factor for developing RA and/or exacerbating ongoing RA [7]. Animal studies also support an association between RA and PD [8–11]. For example, pre-existing PD exacerbates experimental arthritis in a mouse model [11]. Mechanistic studies have proposed that *Porphyromonas gingivalis* induces the production of anti-citrullinated peptide antibodies, which are present in most RA patients, and both *P. gingivalis* and *P. nigrescens* induce Th17 cells, which are known to promote bone erosion via activation of antigen-presenting cells through Toll-like receptor 2 and production of interleukin (IL)-1 and, probably, IL-6 [12]. However, one study found a significant association between RA and PD, but not between RA and the presence of *P. gingivalis* in subgingival biofilms [13]. Thus, the mechanisms underlying the relationship between RA and PD remain unclear.

Numerous plant-derived components have been documented to have anti-inflammatory effects [14–16]. For example, apigenin, which is abundant in common fruits and vegetables, inhibits lipopolysaccharide-induced inflammatory responses in macrophages through multiple mechanisms [14]. *Canavalia gladiate*, also known as the sword bean, is a domesticated plant species native to tropical Asia and Africa, and its fruit has long been used in oriental herbal and folk medicine for treating pus discharge. We previously reported that sword bean extract (SBE) inhibited the growth of *P. gingivalis* and *Fusobacterium nucleatum* and completely suppressed *P. gingivalis-induced* resorption of alveolar bone in an animal model [17]. In a pilot study, we also found that SBE significantly inhibited tumor necrosis factor (TNF)-α and interleukin (IL)-1β production from lipopolysaccharide-stimulated THP-1 (macrophage/monocyte) cells (S1 Fig), suggesting a possible mechanism for the anti-inflammatory effects of SBE.

In the present study, we employed the SKG mouse model of RA to examine the association between RA and ligature-induced PD, and investigated the anti-inflammatory effect of SBE in mice with RA and/or PD.

## Material and Methods

### Animals

4-weeks-old female SKG mice were purchased from Hokudo (Tokyo, Japan) and were maintained at constant temperature in a specific pathogen-free room with a 12-h light-dark cycle. Mice were acclimated for 1 week before experiments. A total of 48 mice were assigned to eight groups of six for the following treatments: (1) controls (no treatment), (2) RA (induced at 6 weeks of age), (3) PD (induced at 10 weeks of age), (4) RA + PD, (5) SBE (added to drinking water starting at 5 weeks of age), (6) PD + SBE, (7) RA + SBE, and (8) RA + PD + SBE. All animals were sacrificed at 13 weeks of age. The experimental design is shown in Fig 1. The study was approved by the Committee of Ethics on Animal Experiments at the Health Sciences University of Hokkaido (approval number 68) and was carried out according to the guidelines for animal experimentation of the Health Science University of Hokkaido.

**Fig 1.**
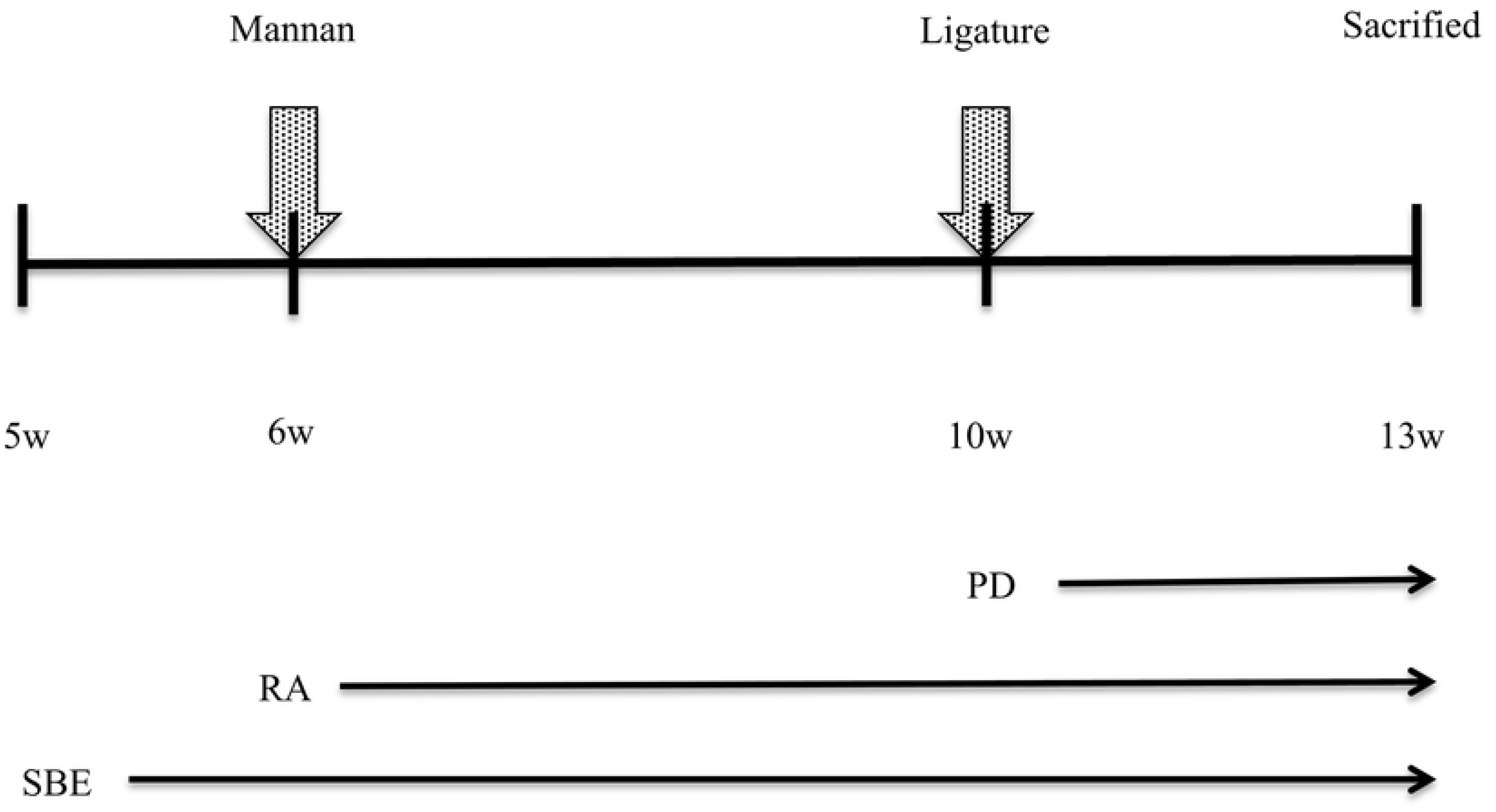
Experimental protocol. The study duration and key time points are indicated. RA and PD were induced at 6 and 10 weeks of age, respectively (See Methods for details). Mice were provided free access to drinking water with or without SBE (2 mg/ml). All animals were sacrificed at 13 weeks of age.

### Preparation of SBE

Sword beans were cultivated in Kagoshima, Japan and SBE was prepared by Yoshitome Co., LTD. (Chiba, Japan). In brief, ground sword beans were mixed with 50% ethanol and filtered to remove the insoluble fraction. The extract was lyophilized to a powder, which was reconstituted in water for use in experiments [17].

### RA induction

Mannan was purchased from Sigma-Aldrich (Tokyo, Japan) and dissolved in phosphate-buffered saline at 100 mg/ml. RA was induced when mice were 6 weeks old by intraperitoneal injection of 200 μl of the mannan solution [18].

### PD induction

At 10 weeks of age, mice were anesthetized with pentobarbital sodium (50 mg/kg) and PD was induced by tying a 5-0 silk ligature (Alfresa Pharma, Osaka, Japan) around the maxillary second molar on both sides (Kyoritsu Seiyaku, Tokyo, Japan). The ligature was left in place until sacrifice 3 weeks later [19].

### Clinical assessment of RA

Paw swelling was assessed and scored by three examiners who were blinded to the treatments at every weeks until the end of the experiment. Each paw was given a visual score (0, no joint swelling; 0.1, swelling of one digit or toe joint; 0.5, mild swelling of the wrist or ankle; and 1.0, severe swelling of the wrist or ankle) and the four scores were summed to give the total score for each mouse [20].

### Morphometric analysis of alveolar bone loss

Using a digital camera system, the buccal surface of the maxillary posterior dentition was recorded in a standardized manner using fixed reference points on the jaws. The surface perimeters of the cement–enamel junction and alveolar bone crest were traced using Adobe Photoshop (Adobe Systems, San Jose, CA, USA), and the software line tool was used to make bone resorption measurements on second molars in each quadrant from the cement–enamel junction to the alveolar bone crest. The area of bone loss (in mm^2^) was printed over each digital image. The areas between the cement–enamel junction and alveolar bone crest were calculated using Image J analysis software (National Institutes of Health, Bethesda, MD, USA) [21].

### Histological analysis

The maxillae and front paws were collected, fixed with 10% buffered formalin solution for 24 h, decalcified in 10% ethylenediaminetetraacetic acid, and then embedded in paraffin wax. Serial sections (periodontal tissue 5 μm, paw 7 μm) were cut, stained with hematoxylin and eosin, and observed for inflammation of the periodontal tissue and synovium. Inflammation of the synovium was scored as follows: 0, no hyperplasia or inflammation; 1, slight hyperplasia with scattered acute inflammation; 2, multiple foci of inflammation with neutrophil predominance; and 3, high levels of inflammatory cell infiltration [10].

### Immunohistochemical (IHC) staining

To detect cathepsin K, serial tissue sections were deparaffinized, rehydrated, and incubated with 3% hydrogen peroxide in methanol for 15 min at room temperature to quench endogenous peroxidase activity. The slides were then incubated with a primary mouse monoclonal antibody against cathepsin k (Kyowa Pharma Chemical) at 4°C for 90 min. A secondary antibody (Simple Stain Mouse Kit, Nichirei Biosciences, Tokyo, Japan) was incubated according to the manufacturer’s instructions. Color development was achieved with 3,3’-diaminobenzidine, which renders positively stained cells brown. Photographs were captured using an Olympus BX-50 upright microscope (Tokyo, Japan).

For IL-17 staining, serial sections were treated in the same manner as described above up to and including the peroxidase quenching step. Antigen retrieval was then carried out by placing the slides in 0.01 M sodium citrate buffer (pH 6.0) at 95°C for 30 min. The slides were incubated with a primary rabbit polyclonal antibody against IL-17 (ab 19046 Abcam, Cambridge, UK) at 4°C overnight. Color development and imaging of the slides was performed as described for cathepsin K.

### Serum C-reactive protein (CRP) quantification

Blood samples were collected from mice before sacrifice by cardiac puncture, left at room temperature for 2 h to clot, and then centrifuged at 1000 ×g for 20 min. The sera were collected and stored at −80°C until quantification of CRP levels using a Mouse CRP ELISA Kit (Life Diagnostics, West Chester, PA, USA).

### Statistical analysis

Data are presented as the mean ± standard error (SEM) and were analyzed using JMP 8 software. Group means were compared using one-way analysis of variance (ANOVA) with Tukey’s post hoc test. *P* < 0.05 was considered statistically significant.

## Results

### Clinical assessment

Mice in the RA and RA + PD groups had more severe paw redness and swelling than mice in the other groups (Fig 2A). By the end of the experiment (13 weeks of age), both symptoms were more severe in the RA + PD mice than in the RA mice. However, SBE treatment reduced the paw redness and swelling in the RA + PD mice at 13 weeks (Fig 2B).

**Fig 2.**
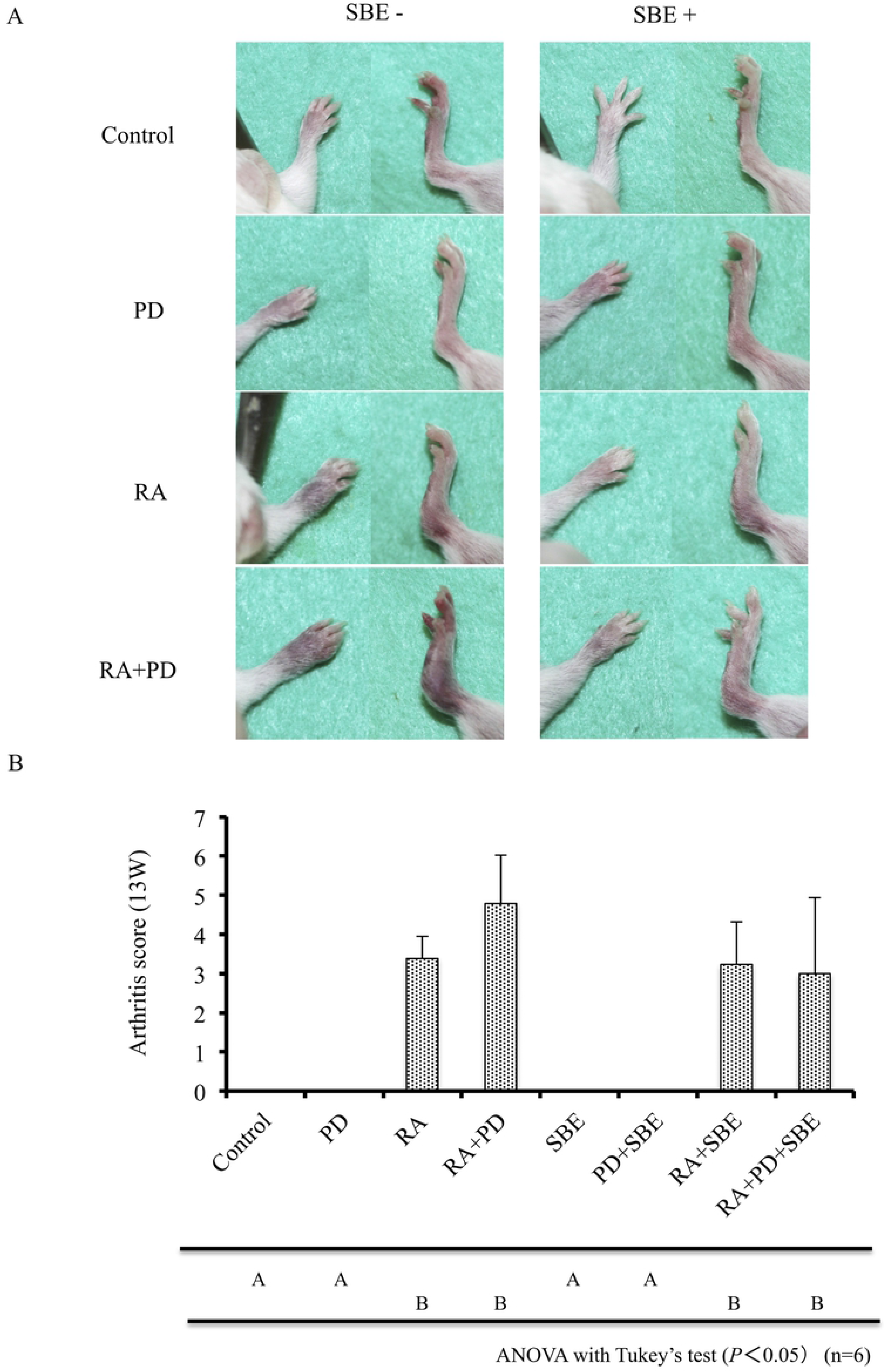
Clinical assessment of RA. A. Clinical appearance of the front and rear paws. B. Average paw arthritis score at 13 weeks of age (mean ± SD, n=6). Data were analyzed by one-way ANOVA with Tukey’s test. Letters indicate statistical significance (*P* < 0.05). RA, rheumatoid arthritis; PD, periodontal disease; SBE, sword bean extract.

### Alveolar bone loss

Morphometric analysis at 13 weeks showed that alveolar bone loss was significantly higher in the mice in the PD and RA + PD groups compared with any other group, and was significantly higher in the RA + PD than in the PD mice. SBE treatment significantly reduced the alveolar bone loss in all mice with PD (Figs 3A and B).

**Figure 3.**
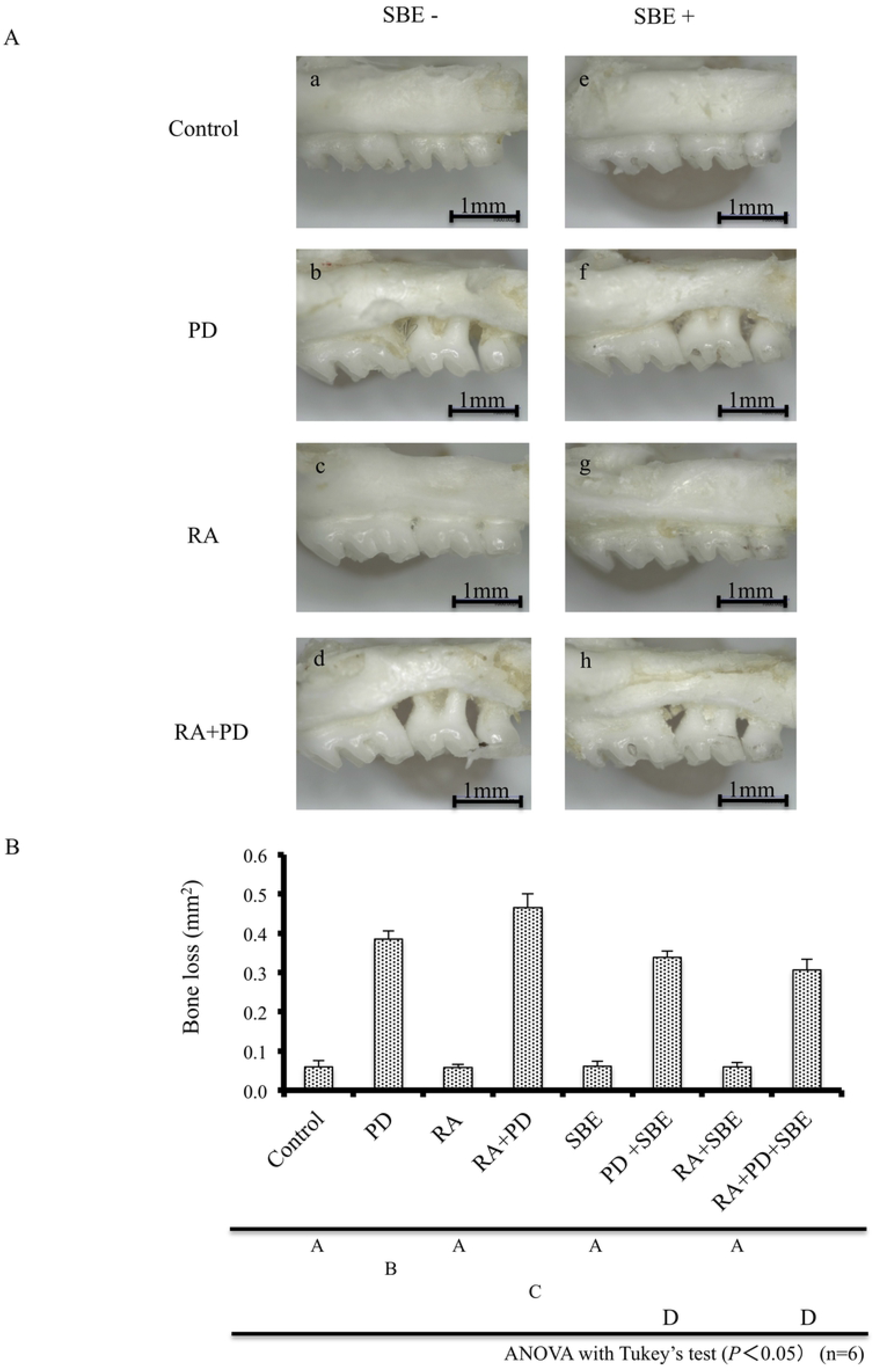
Morphometric analysis of alveolar bone loss. A. Representative photographs illustrating the morphometric findings. B. Average alveolar bone loss (mm^2^) at 13 weeks of age (mean ± SEM, n=6). Data were analyzed by one-way ANOVA with Tukey’s test. Letters indicate statistical significance (*P* < 0.05). RA, rheumatoid arthritis; PD, periodontal disease; SBE, sword bean extract.

### Histological analysis

Bone resorption, infiltration of inflammatory cells, and proliferation of synovial cells were not observed in the radicular carpal joints of the control mice, PD mice, SBE mice, or PD + SBE mice (Figs 4Aa, a1, b, b1, e, e1, f and f1). However, bone resorption and pathological proliferation of the synovium accompanied by infiltration of inflammatory cells were observed in the RA mice, RA + PD mice, RA + SBE mice, and RA + PD + SBE mice (Figs 4Ac, c1, g, g1, h and h1). This phenotype was most prominent in the RA + PD group (Figs 4Ad and d1).

**Fig 4.**
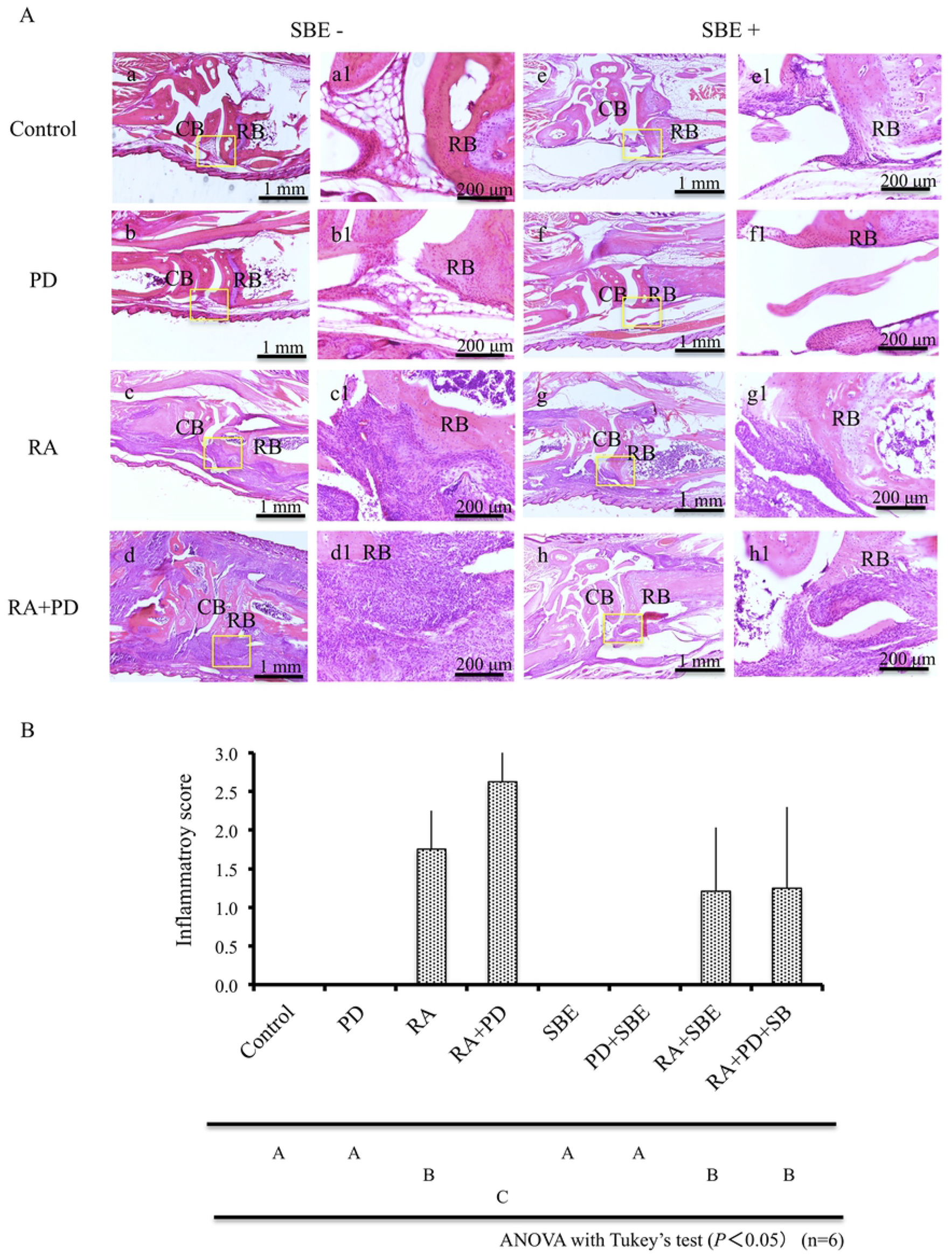
Histological analysis of RA. A. Histological appearance of front paw joints at ×4 magnification (a, b, c, d, e, f, g, h) and ×20 magnification (a1, b1, c1, d1, e1, f1, g1, h1). Inflammation scores are shown as the mean ± SD (n=6). Data were analyzed by ANOVA with Tukey’s test. Letters indicate statistical significance (*P* < 0.05). RA, rheumatoid arthritis; PD, periodontal disease; SBE, sword bean extract; RB, radial bone; CB, carpal bone.

Inflammation scores were significantly higher for the RA + PD mice than the RA mice, and significantly lower for the RA + PD + SBE mice than for the RA + PD mice (Fig 4B). There was no significant difference between the inflammation scores for the RA mice and the RA + SBE mice, although the scores tended to be lower for the SBE-treated group (Fig 4B).

### Histological analysis

Mice in the control, RA, SBE, and RA + SBE groups did not exhibit either bone resorption in the interproximal parts of the maxillary first molar and the second molar, or inflammatory cells in the gingiva and periodontal ligament around the alveolar bone (Figs 5 a, a1, c, c1, e, e1, g and g1). In contrast, bone resorption and inflammatory changes were seen in mice in both the PD and RA + PD groups (Figs 5 b, b1, d and d1). While the PD + SBE and RA + PD + SBE groups showed bone resorption, inflammatory cells were virtually undetectable in the periodontal tissues (Figs 5 f, f1, h and h1).

**Fig 5.**
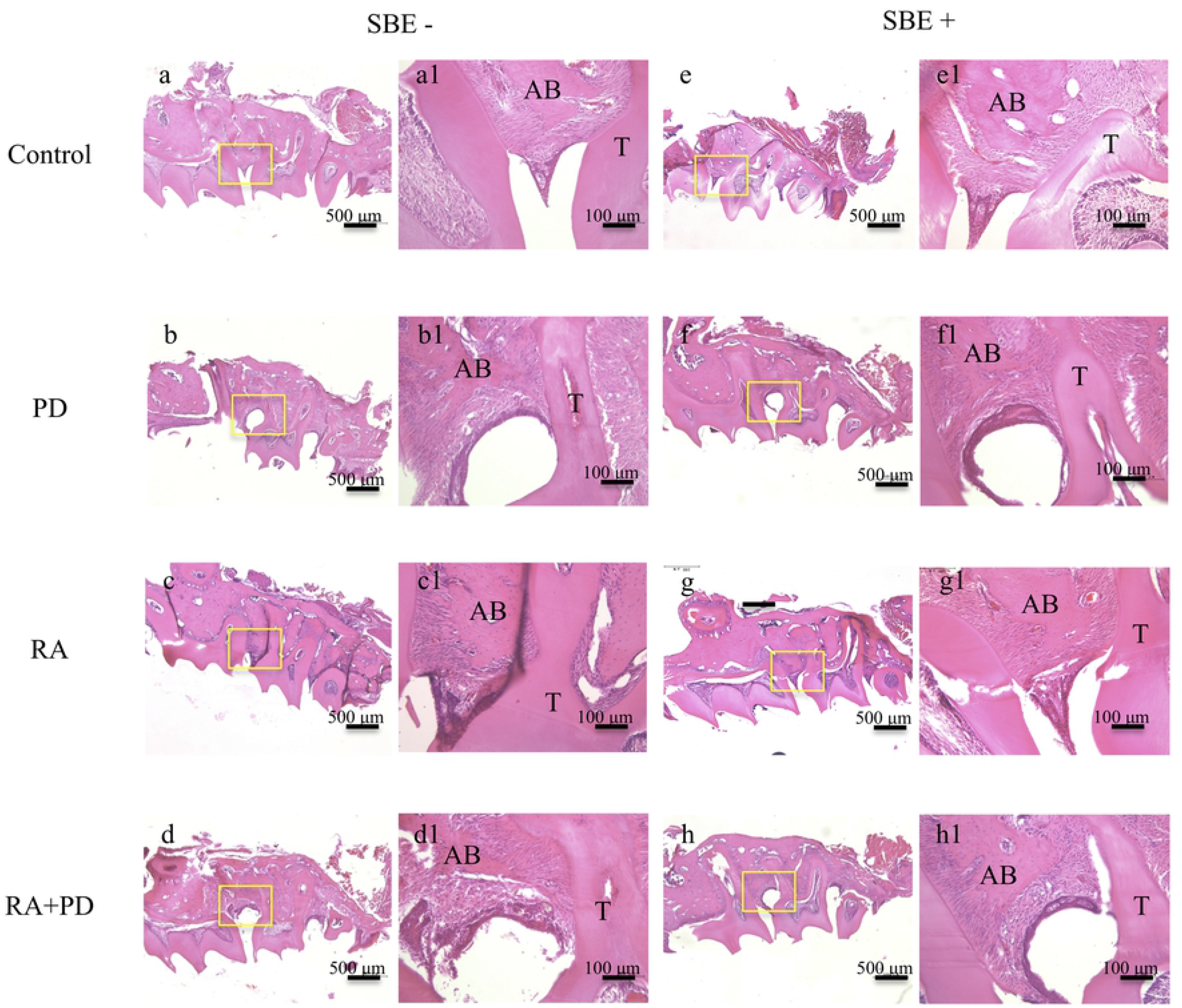
Histological analysis of PD. Histological appearance of periodontal tissue at ×8 magnification (a, b, c, d, e, f, g, h) and ×40 magnification (a1, b1, c1, d1, e1, f1, g1, h1). RA, rheumatoid arthritis; PD, periodontal disease; SBE, sword bean extract. T, teeth; AB, alveolar bone,

### IHC analysis of IL-17

Although very few IL-17-positive cells were detected around the radial bone of mice in the control, PD, SBE, and PD + SBE groups (Figs 6a, b, e and f), they were present in the RA, RA + SBE, and RA + PD + SBE groups (Figs 6c, g and h). IL-17-positive cells were particularly abundant in mice in the RA + PD group (Fig 6d).

**Figure 6.**
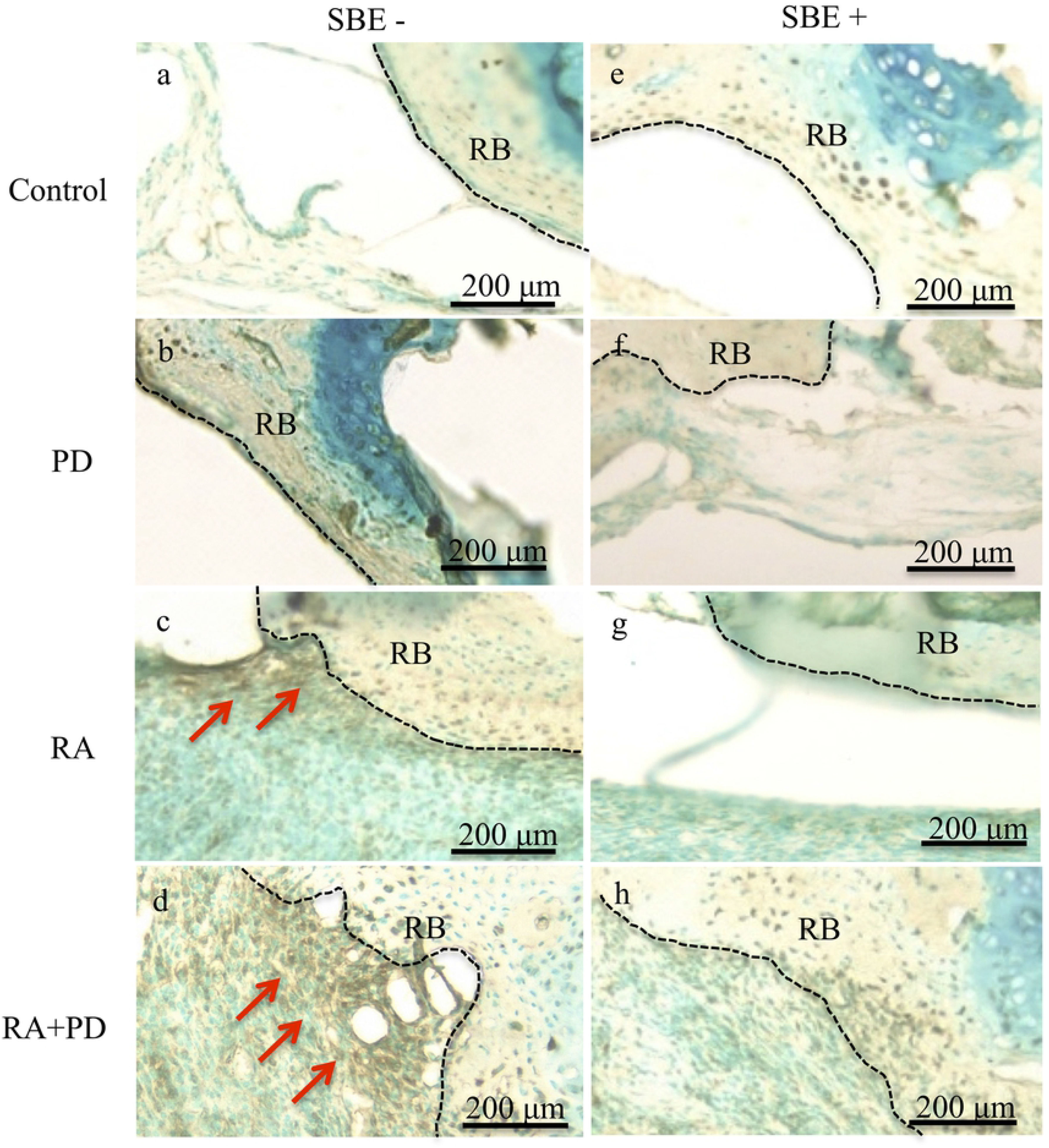
IHC analysis of IL-17-expressing cells in mice with RA. Representative joint tissue sections from SKG mice showing IHC staining of IL-17 at ×20 magnification. Red arrows indicate IL-17-positive cells. RA, rheumatoid arthritis; PD, periodontal disease; SBE, sword bean extract; RB, radial bone; CB, carpal bone.

### IHC analysis of Cathepsin K

IHC analysis of the front paws indicated that cathepsin K-positive cells were significantly more abundant in the RA + PD mice compared with the RA mice, but significantly scarcer in RA + PD + SBE mice than in RA + PD mice. Although fewer cathepsin K-positive cells were detected in the paws of RA + SBE mice than in RA mice, the difference did not reach the level of statistical significance (Figs 7A and B).

**Figure 7.**
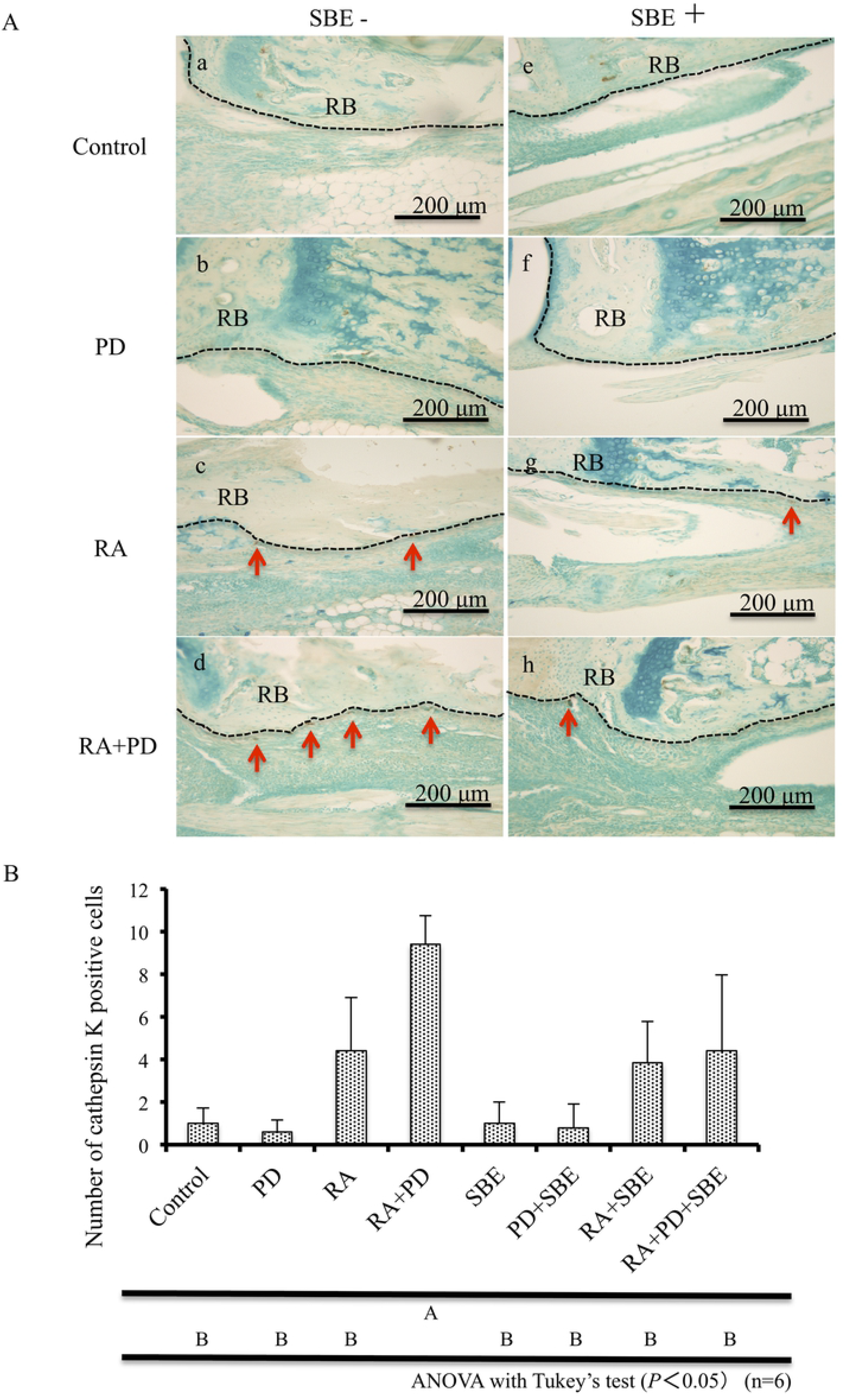
IHC analysis of Cathepsin K-positive cells in mice with RA. A. Representative bone sections from SKG mice showing IHC staining of cathepsin K at ×20 magnification. Red arrows indicate cathepsin K-positive cells. B. Enumeration of cathepsin K-positive cells surrounding the radial bone (mean ± SD, n=6). Data were analyzed by one-way ANOVA and Tukey’s test. Letters indicate statistical significance (*P* < 0.05). RA, rheumatoid arthritis; PD, periodontal disease; SBE, sword bean extract; RB, radial bone; CB, carpal bone.

### Serum CRP levels

CRP concentrations were significantly higher in serum samples from the RA and RA + PD mice than the control or PD mice and significantly higher in the samples from the RA + PD mice compared with the RA mice. SBE treatment reduced the CRP levels in RA + PD mice and RA mice, but the difference was statistically significant only for the RA + PD groups (Fig 8).

**Figure 8.**
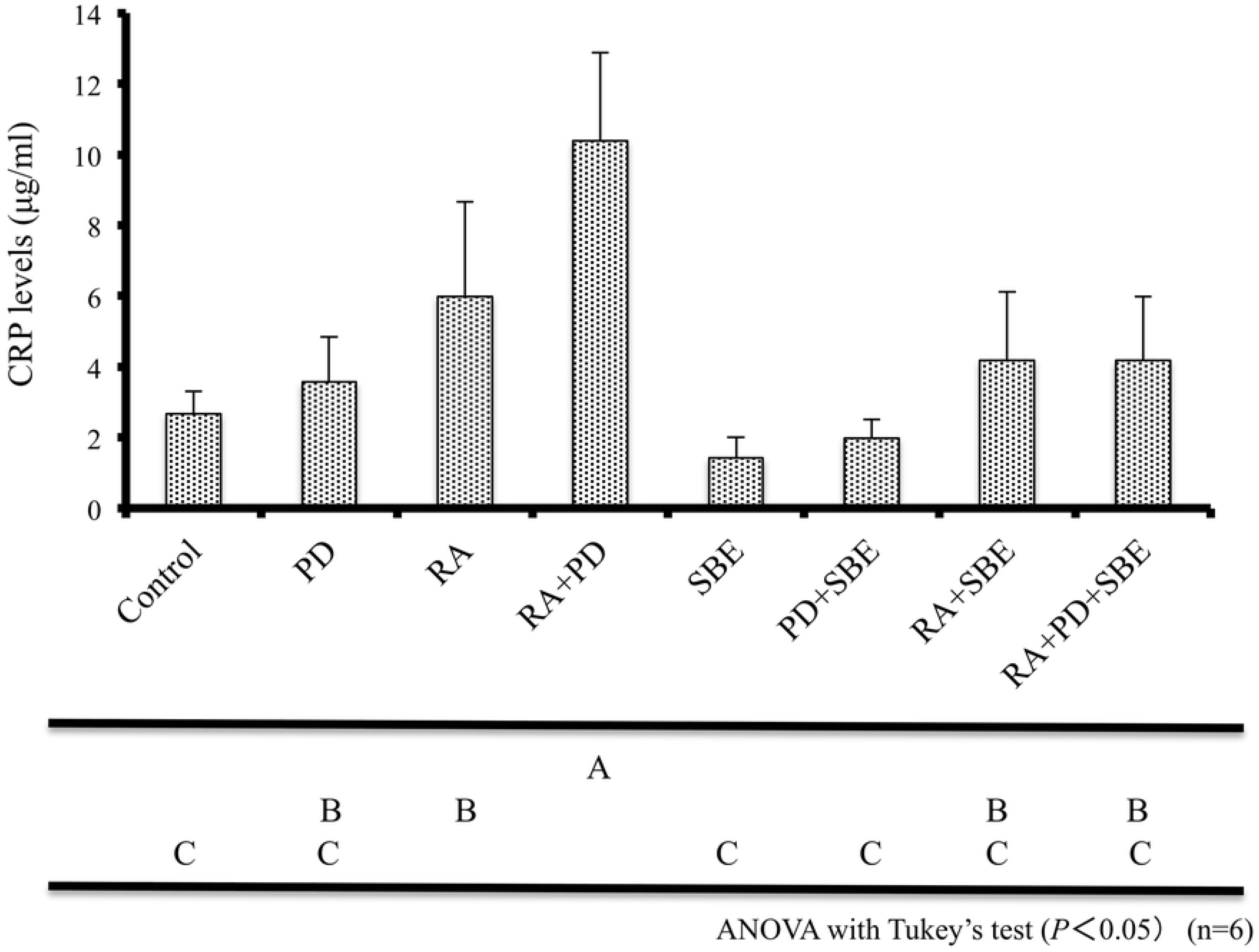
Serum CRP levels. Average CRP levels measured by ELISA. data are shown as the mean ± SD. Data were analyzed by one-way ANOVA and Tukey’s test. Letters indicate statistical significance (*P* < 0.05). RA, rheumatoid arthritis; PD, periodontal disease; SBE, sword bean extract.

## Discussion

Several reports have demonstrated an association between RA and PD in humans as well as experimental animals. In the present study, we evaluated the association between these diseases in SKG mice. We found that concomitant RA and PD exacerbated both PD-associated alveolar bone resorption and RA-associated joint inflammation. These findings are consistent with the results of a previous study in which PD was induced in a similar manner in a rat experimental arthritis model [22]. Among the various reports on the association between RA and PD, several have implicated periodontal pathogenic bacteria such as *P. gingivalis, Treponema denticola*, and *Tannerella forsythia*, the major PD-related species [23]. Since *P. gingivalis* can produce peptidyl arginine deiminase, which citrullinates proteins, it has been suggested to play an important role in the development of RA [24]. For example, *P. gingivalis* infection exacerbates arthritis in a collagen antibody-induced mouse model [11] and in another SKG mouse model of RA [25]. However, PD, but not *P. gingivalis* colonization, was associated with arthritis activity in treatment-naïve arthritis patients [13]. We believe that *P. gingivalis* is unlikely to be involved in the alveolar bone loss reported in the current study because we induced PD by ligation. Indeed, our finding that alveolar bone resorption and joint inflammation scores were significantly higher in mice with RA and PD than in mice with either disease alone suggests that they interact to induce inflammation without bacterial infection. In the present study, SBE suppressed bone resorption, periodontal tissue inflammation, and joint inflammation in mice with concomitant RA and PD. The effect of SBE on periodontal tissues is consistent with our previous study in which we examined a *P. gingivalis* infection model of PD in rats [17]. However, the previous study suggested that SBE acted by inhibiting gingipain activity, which is not a factor in the current study. It has been reported that levels of cytokines involved in bone resorption, such as IL-1β, are increased in the periodontal tissue of mice with ligature-induced PD [26], and that both IL-1β and IL-6 are increased in the serum of such mice [22, 26, 27]. In the ligature-induced PD model, alveolar bone resorption was suppressed by antibacterial treatment, suggesting that inflammation was enhanced by the accumulation of plaque in the oral cavity [19]. In the present study, SBE suppressed the inflammatory response and bone resorption in periodontal tissues, as shown by IHC. We speculate that SBE may act as an antimicrobial agent, as suggested by earlier studies, or as an anti-inflammatory agent, as supported by our pilot study (S1 Fig).

In the present study, IL-17-producing cells and cathepsin K-positive osteoclasts were higher in the radial and carpal bones of RA + PD mice compared with RA mice. IL-17 is an essential mediator of cartilage and bone destruction [28–30] and its production is promoted by inflammatory cytokines such as IL-1β and IL-6 [12]. IL-17 also enhances the differentiation of osteoclasts by upregulation of RANKL (receptor activator of nuclear factor kappa-B ligand) and TNF-α [28, 31]. These observations suggest that IL-17 upregulation and in duction of osteoclast differentiation and activation may have contributed to the bone resorption observed in the present study. Moreover, SBE-mediated suppression of the periodontal inflammatory response may also have dampened the RA symptoms. Although we do not know the mechanism by which SBE inhibits IL-17 production or osteoclast differentiation, our pilot study strongly suggests that SBE inhibits proinflammatory cytokine production.

Smith et al. reported that IL-6 induces the production of CRP by the human liver cancer cell line, HepG2 [32]. Serum IL-6 and CRP concentrations are positively correlated in PD patients and are higher in this population than in healthy subjects [33]. In the present study, we found that serum CRP levels were higher in mice with RA and PD compared with mice with RA or PD alone. One possibility is that CRP is induced by inflammatory cytokines, such as IL-6, that are produced locally in the periodontal tissue but then circulate systemically. Serum CRP concentrations were reported to decrease in patients with PD and RA after periodontal therapy [34]. In our study, CRP levels were reduced in the RA + PD mice by SBE treatment, supporting a possible connection between suppression of inflammatory cytokine production and serum CRP concentrations.

We do not yet know which SBE components are responsible for the anti-inflammatory effects observed here. The composition of sword beans is thought to be similar to that of edible beans, but a detailed analysis of SBE has not yet been conducted. Genistein, a dietary polyphenol primarily found in soy products, is known to protect against inflammatory periodontal damage by regulating autophagy induction and inhibiting osteoclast activation, inflammatory mediator production, and mitochondrial oxidative damage [35]. Similarly, p-coumaric acid, which is abundant in fruits and vegetables, has been shown to improve joint pathology in an adjuvant-induced rat arthritis model [36]. Previous studies by our laboratory showed that SBE contains canavanine, which inhibits the activity of Kgp and Rgp in vitro. However, canavanine at concentrations contained in SBE, is able to suppress alveolar bone resorption in vivo without suppressing the activity of Kgp and Rgp [17]. Suganuma et al. [37] have demonstrated that epigallocatechin, which is contained in green tea, has synergistic anti-cancer effects in combination with gallate and (-)-epicatechin Thus, synergism between various components of SBE may be responsible for its anti-inflammatory effects. In the future, we plan to investigate possible applications for SBE to prevent or treat inflammatory diseases such as RA and PD. It will also be important to identify the constituents of SBE to determine the mechanism of its anti-inflammatory effects.

## Conclusion

In this study, we found that concomitant RA and PD exacerbates the destruction of periodontal and joint tissue in a mouse model, suggesting an association between the two diseases. Alveolar bone resorption and joint inflammation were both significantly suppressed by treatment with SBE. Our preliminary data suggest that SBE inhibits inflammatory cytokine production, providing a possible mechanism by which it may suppress the interaction between RA and PD in vivo.

## Acknowledgments

There is none.

## Supporting information

**S1 Fig.** Sword Beans Extract (SBE) significantly inhibited TNF-α and IL-1β production from LPS-stimulated THP-1 cells. Data were analyzed by one-way ANOVA with Tukey’s test. Letters indicate statistical significance (*P* < 0.05).

